# Ensemble Machine Learning to “Boost” Ubiquitination-sites Prediction

**DOI:** 10.1101/2022.09.11.507485

**Authors:** Xiaoye Mo, Xia Jiang

## Abstract

Ubiquitination-site prediction is an important task because ubiquitination is a critical regulatory function for many biological processes such as proteasome degradation, DNA repair and transcription, signal transduction, endocytoses, and sorting. However, the highly dynamic and reversible nature of ubiquitination makes it difficult to experimentally identify specific ubiquitination sites. In this paper, we explore the possibility of improving the prediction of ubiquitination sites using ensemble machine learning methods including Random Forrest (RF), Adaptive Boosting (ADB), Gradient Boosting (GB), and eXtreme Gradient Boosting (XGB). By doing grid search with the four ensemble methods and six comparison non-ensemble learning methods including Naïve Base (NB), Logistic Regression (LR), Decision Trees (DT), Support Vector Machine (SVM), LASSO, and K-Nearest Neighbor (KNN), we find that all the four ensemble methods significantly outperform one or more non-ensemble methods included in this study. XGB outperforms three out of the six non-ensemble methods that we included; ADB and RF both outperform two of the six non-ensemble methods; GB outperforms one non-ensemble method. Comparing the four ensemble methods among themselves. GB performs the worst; XGB and ADB are very comparable in terms of prediction, but ADB beats XGB by far in terms of both the unit model training time and total running time. Both XGB and ADB tend to do better than RF in terms of prediction, but RF has the shortest unit model training time out of the three. In addition, we notice that ADB tends to outperform XGB when dealing with small-scale datasets, and RF can outperform either ADB or XGB when data are less balanced. Interestingly, we find that SVM, LR, and LASSO, three of the six non-ensemble methods included, perform comparably with all the ensemble methods. Based on this study, ensemble learning is a promising approach to ignificantly improving ubiquitination-site prediction using protein segment data.

## BACKGROUND

Ensemble learning is a technique that learns multiple models from the same dataset and then combines them to form an optimal learner, which is anticipated to have better prediction performance. Supervised learning provides algorithms to perform a searching task with limited hypothesis space to find an optimal hypothesis which will manage the prediction task successfully with a specific problem [1]. However, it can be challenging to find a suitable hypothesis for a particular problem, even when the hypothesis space is well designed. The ensemble method in machine learning, which combines different hypotheses to hopefully construct a much better hypothesis, has been discussed for decades. With multiple models, it can reduce the classification error rate in a classification task [2].

Empirically, ensemble learning is expected to provide better results when a substantial diversity is found among models, which sometimes may cause over-fitting or under-fitting [3, 4]. With this experience, researchers tried to develop many ensemble learning methods which could take advantage of the diversity among different models and then combine them together [3, 5, 6]. Although it may not be intuitive, stochastic algorithms (e.g., random decision trees) could be used to generate stronger ensemble models than intentional algorithms (e.g., entropy-reducing decision trees) [7]. Moreover, using a variety of powerful learning algorithms has already been shown to be more effective than using techniques that try to simplify the model to promote diversity [8].

Although there can be many ways to implement ensemble learning, we focused on two approaches which are widely discussed and applied in practice [9–12]: bootstrap aggregating and boosting. Bootstrap aggregation, also called bagging (from bootstrap aggregating), is usually applied to decision tree methods, but can also be applied to any other method [13–15]. As implied in its name, bootstrap and aggregation are two key components of bagging. Bootstrap is a sampling method. Given a standard training set *D* of size *n*, bagging generates *m* new sub-training sets *D_i_*, each of size *n*′, by sampling from *D* randomly with replacement. By sampling with replacement, some observations may be repeated in each *D_i_*), but some observations may not appear in any subset *D_i_*. When sampling uniformly, if *n*′ = *n*, for large *n*, the set *D_i_* is expected to have the fraction (1 – 1/*e*) (≈ 63.2%) of the unique examples of the original set *D*, the rest being duplicates [16]. After the bootstrapped sampling, several sub-datasets are created. Each sub-dataset will be used by a machine learning method to train a model, which will result in multiple models being trained by using different sub-datasets. Since each bootstrap set is randomly generated, the sets of the sub-datasets are expected to be diversified, and therefore, the individual models in the ensemble are expected to represent a different aspect of the original data. Due to this, when combining all sub-models together, the ensemble is expected to represent aspects of the original data. Finally, the ensemble will make predictions through simple statistics such as voting. Figure 1(A) on the left illustrates the general components and procedures of bagging.

**Figure 1.**
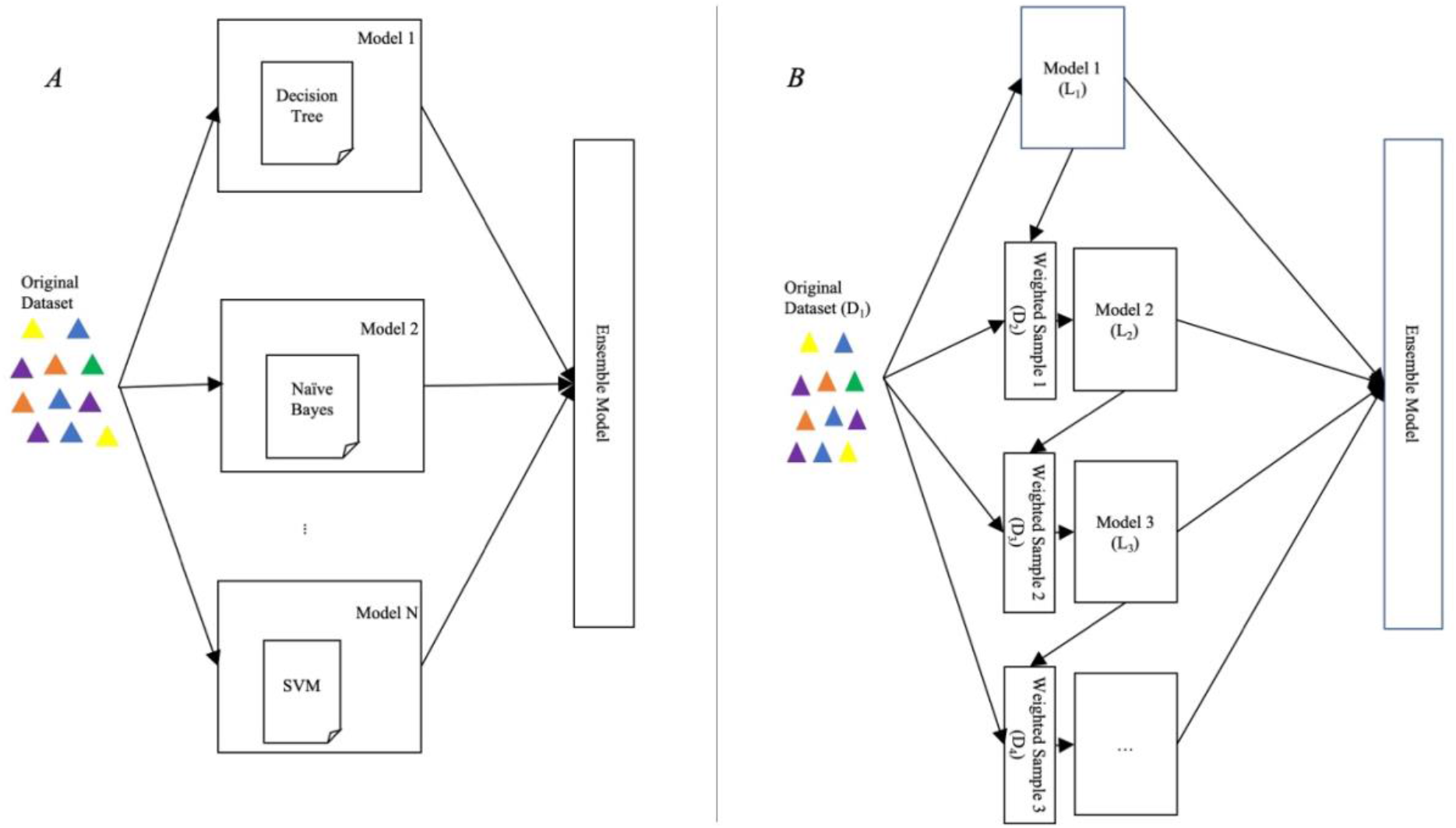
An illustration of ensemble methods (A) Bagging (B) Boosting

Boosting is the third main approach of ensemble learning, which involves incrementally building the ensemble and training new model instances by focusing on wrong results made by previous trained models [17–19]. Boosting comes from the famous question from Kearns and Valiant: “Can a set of weak learners create a single strong learner?” [20]. In 1990, Robert offered an answer to that question, which significantly influenced machine learning, and led to the development of boosting [19]. Later, researchers have proved that boosting can have better accuracy than bagging in some cases, but it may not handle the over-fitting issue as well [21]. Figure 1(B) on the right shows the general idea of boosting: the same weight is given to each training data *D*_1_ at the beginning. The base learner *L*_1_ will learn Model1 from *D*_1_. Then, the wrongly classified instances by Model1 will be given a larger weight than the correct ones. The second base learner *L*_2_ will learn Model 2 from the weighted data *D*_2_, and so on and so forth. The voting algorithm is applied in the final round to produce the results.

Ubiquitination, also called ubiquitylation is an enzymatic and post-translational modification process in which ubiquitin, a small regulatory protein, is attached to substrate proteins [22, 23] During the ubiquitination process, ubiquitin interacts with lysine (K) residues on protein substrates through three steps: ubiquitin-activating enzymes (E1s), ubiquitin-conjugating enzymes (E2s), and ubiquitin ligases (E3s) [22–24]. It needs to be mentioned that binding can be a single ubiquitin or a ubiquitin chain. It has been found that many regulatory functions of ubiquitination, such as proteasome degradation, DNA repair and transcription, signal transduction, endocytosis, and sorting, are important protein regulatory functions in biological processes [22–25].

Because ubiquitination plays an important regulatory role, extensive research has been conducted to further decipher the mechanism of ubiquitination and other regulatory roles at the molecular level. One of the initial and challenging steps in gaining a greater understanding of ubiquitination is to identify ubiquitination sites. To purify ubiquitinated proteins to determine ubiquitination sites, researchers have used different types of experimental methods, such as high-throughput mass spectrometry (MS) techniques [26–29], ubiquitin antibodies and ubiquitin-binding proteins [29, 30],and combined liquid chromatography and mass spectrometry [31]. However, experiments to purify ubiquitinated proteins are time-consuming, expensive, and labor-intensive because the ubiquitination process is dynamic, rapid, and reversible [24, 32, 33]. To reduce experimental costs and increase the effectiveness and efficiency of ubiquitination site identification, computational (in silico) methods based on informatics techniques have been introduced and developed for predicting ubiquitination sites based on prior knowledge of protein sequences [24, 25, 32, 33].

Previous research applied some basic machine learning methods to predict ubiquitination-site with PhysicoChemical Property (PCP) datasets [34]. As we know, a protein is a biological molecule that consists of one or more long chains of amino acid residues; PCP datasets are generated from protein sequences and then are processed by several steps, including segment extraction, creating AA-PCP matrix, and averaging [34]. Although the study demonstrated the potential of conducting this challenging task by taking the machine learning approach, results from the base machine learning methods were not completely satisfactory. Since ensemble methods are proved to do better than base learners in many cases, we conjectured that they are potentially powerful in predicting ubiquitination-sites. In this research, we considered four widely used state of art ensemble methods including Random Forrest (RF), Adaptive Boosting (ADB), Gradient Boosting (GB), and eXtreme Gradient Boosting (XGB). To verify our conjecture, we conducted ubiquitination-site prediction by learning prediction models using these methods from previously published PCP datasets [34].

## METHODS

As described above, we did the experiments with two widely used types of ensemble methods, Bagging and Boosting. RF is one of the most famous Bagging approaches of ensemble learning [35]. RF was published in 2001 by Breiman, which utilizes Bagging to generate different subsets for entire training to build individual decision trees [35].Then, RF has been applied into different fields of research, including chemical, power transmission, ecology, etc. [14, 36, 37]. ADB, GB, and XGB are three Boosting approaches. AdaBoost was purposed by Freund and Schapire around 1996 [18, 38]. In 2001, Friedman proposed the GB method, opening the door to Boosting [39].XGB, a new development based on GB, was proposed by Chen and He in 2014 [40]. Unlike RF, the other three methods are Boosting algorithms, and the main idea of boosting is to build models sequentially based on previous trained models [41].

Since ensemble methods were said to improve the performance of prediction models in many cases, we tried to apply them on the ubiquitination-site prediction task. Hyperparameter tuning with grid search is a state-of-the-art approach to improve prediction performance. In this study, we optimized 4 ensemble learning models by performing hyperparameter tuning via grid search. We next describe the six PCP datasets we used. We will then describe each of the four existing ensemble methods we used, including RF, ADB, GB, and XGB. We will also describe the grid search we conducted, the hyperparameter values we used in the grid search for each of these methods, and the evaluation metrics we used.

### Datasets

The six (PCP) datasets were curated and published in 2016 by Cai and Jiang [34]. As shown in Table 1, each dataset contains 531 features. These 6 datasets can be divided by 2 types. PCP 1-3 are balanced datasets, meaning each dataset contains the same number of positive and negative cases, while PCP 4-6 are imbalanced datasets with unequal numbers of positive and negative cases. Also, PCP 1 and 6 are small-scale datasets with less than a thousand cases each, and PCP 2-5 are large-scale datasets which contain thousands of data points each.

**Table 1.** Case counts of the datasets.

| **Dataset** | **Total # of cases** | **# Positive cases** | **# Negative cases** | **# Columns** |
| --- | --- | --- | --- | --- |
| PCP-1 | 300 | 150 | 150 | 531 |
| PCP-2 | 6838 | 3419 | 3419 | 531 |
| PCP-3 | 12236 | 6118 | 6118 | 531 |
| PCP-4 | 4608 | 363 | 4345 | 531 |
| PCP-5 | 3651 | 131 | 3520 | 531 |
| PCP-6 | 676 | 37 | 639 | 531 |

### Ensemble Methods

Ensemble learning is a combination machine learning method that combines the prediction results from multiple models to have an overall good performance. Among the three most popular types of ensemble learning, there are some shared points. First, ensemble learning models all use individual models to train, which means they combine several trained models, unlike traditional machine learning models which are only trained once. However, all “small” prediction models may use different combinations to build the final ensemble model [42]. Bagging, stacking, and boosting are the three most popular methods of combination. Random Forest is one of the most popular bagging methods [35], AdaBoosting, Gradient Boosting, and Extreme Gradient Boosting are widely used boosting algorithms [39, 43, 44]. Second, while traditional machine learning methods usually need constraints on a dataset (for example, a balanced dataset), ensemble learning can handle imbalance problems, with a well-designed ensemble model, it can perform well [45]. In this study, we performed ensemble learning with the methods mentioned above, to analyze how they work with ubiquitination-site prediction.

### Ensemble Machine Learning Methods

We compared the performance of a set of ensemble machine-learning methods, including Random Forest (RF), Adaptive Boosting (ADB), Gradient Boosting (GB), and eXtreme Gradient Boosting (XGB). Like most machine learning methods, these methods have hyperparameters that can be tuned to provide optimal performance. We conducted a grid search for each method for 6 PCP datasets. We conducted 5-fold cross-validation for each set of hyperparameter values and measured the performance by the AUC of ROC (Receiver Operating Characteristic). Below, we provide a summary of the hyperparameters and their values that we tested for each of these 4 ensemble methods. All the test values are shown in Table 2.

**Table 2.**
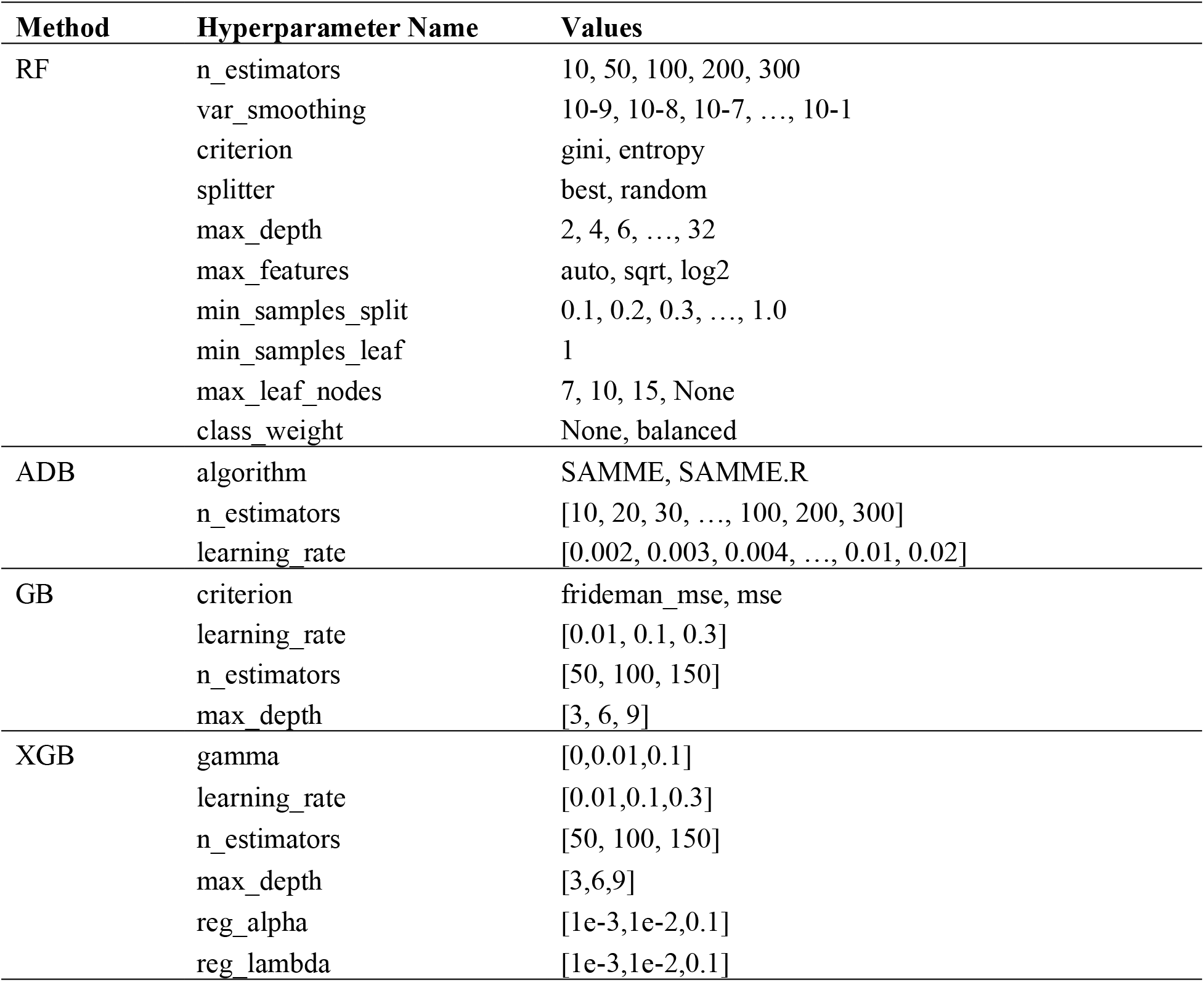
Machine learning hyperparameters and values

| Method | Hyperparameter Name | Values |
| --- | --- | --- |
| RF | n_estimators | 10, 50, 100, 200, 300 |
|  | var_smoothing | 10-9, 10-8, 10-7, ..., 10-1 |
|  | criterion | gini, entropy |
|  | splitter | best, random |
|  | max_depth | 2, 4, 6, ..., 32 |
|  | max_features | auto, sqrt, log2 |
|  | min_samples_split | 0.1, 0.2, 0.3, ..., 1.0 |
|  | min_samples_leaf | 1 |
|  | max_leaf_nodes | 7, 10, 15, None |
|  | class_weight | None, balanced |
| ADB | algorithm | SAMME, SAMME.R |
|  | n_estimators | [10, 20, 30, ..., 100, 200, 300] |
|  | learning_rate | [0.002, 0.003, 0.004, ..., 0.01, 0.02] |
| GB | criterion | frideman_mse, mse |
|  | learning_rate | [0.01, 0.1, 0.3] |
|  | n_estimators | [50, 100, 150] |
|  | max_depth | [3, 6, 9] |
| XGB | gamma | [0,0.01,0.1] |
|  | learning_rate | [0.01,0.1,0.3] |
|  | n_estimators | [50, 100, 150] |
|  | max_depth | [3,6,9] |
|  | reg_alpha | [1e-3,1e-2,0.1] |
|  | reg_lambda | [1e-3,1e-2,0.1] |

**RF** [7, 11, 35, 36, 46] is a typical model of bagging in ensemble learning, the trainer will randomly select a certain amount of sample data and create corresponding decision trees to form a random forest [11]. An advantage of Random Forest is that the independent character of each decision tree tends to reduce overfitting [36, 46]. *n_estimators*: The number of decision trees in the random forest. We tested values 10, 50, 100, 200, 300. The parameter *max_depth* indicates how deep the tree can be; the deeper the tree, the more splits it will have, and thus capturing more information about the data [7, 11, 35]. We fit a decision tree with depths ranging from 2 to 32. The parameter *min_samples_split* represents the minimum number of samples required to split an internal node. The values we tested in our grid search were 0.1, 0.2, 0.3, 0.4, 0.6, 0.8, 0.9 and 1. *max_features* indicated the maximum number of features allowed when building a decision tree [46, 49, 50]; we tested all values under ‘none’, ‘log2’ and ‘sqrt’. The parameter *max_leaf_nodes* controls the maximum number of leaf nodes of each decision tree [7, 11, 35],and we tested values 7, 10, 15, and none. The parameters *max_depth* and *max_leaf_nodes* are important in controlling overfitting [36]. The function *criterion* was used to measure the quality of a split [7, 11, 35];we tested with values ‘gini’ and ‘entropy’.

**ADB** [11, 18, 35, 47, 48] is a typical model of boosting in ensemble learning [11, 35]. Unlike the Random Forest model, where each decision tree is independent, Adaboost is a classifier with a cascade structure, which means the next learner is based on the result of the previous weak learner [18, 35]. During the learning process, if the current sample is classified incorrectly, the degree of difficulty of the sample will increase to make the next learner focus on the difficult part on which the previous model performed poorly [47, 48]. *n_estimators*: The number of weak learners. A model to overfit for large values of *n_estimators*; the values of *n_estimators* we tested include 10, 20, …, 100, 200, and 300. *learning_rate*: this is used to shrink the contribution of each classifier; we tested values include 0.002, 0.003, 0.004, …, 0.01, and 0.02.

GB [39, 41, 49–52] is an ensemble of weak predictions models [39]. Unlike the bagging methods, boosting builds the mode in sequentially [52]. Boosting fits base learners additively to have better performance than random [41, 50, 51]. *learning_rate*, one of the most important hyper-parameters, we tested 0.01, 0.1, 0.3. The number of boosting stages to perform, *n_estimator*, we tried 50, 100, and 150 to test. The values of *max_depth* we tested include 3, 6, and 9. *criterion* which is a function to measure the split, and here we tested two different criterions, ‘friedman_mse’ and ‘mse’.

XGB [40, 43, 52, 53]is another common approach of boosting in ensemble learning. Unlike ADB, it uses gradient boosting [52].The XGB classifier is based on the difference between true and predicted values to improve model performance [40, 53]. *gamma*: this is a pseudo-regularization hyperparameter in gradient boosting; gamma affects pruning to control overfitting problems. *gamma* values we tested were 0, 0.01, and 0.1. *alpha* and *lambda* are both regularization hyperparameters which can help control overfitting [40]. The values we tested for each of them were 1e-3, 1e-2, and 1e-1. *max_depth* is the maximum depth of the individual regression estimators. The values of *max_depth* we tested were 3, 6, and 9. The *learning_rate* values we tested were 0, 0.01, 0.1, and 0.3.

### Grid Search

Machine learning algorithms have been widely used in different types of tasks in the real world. To have machine learning models perform well in a specific task, it often needs a tuning procedure with which we can find a good set of hyperparameters values [54]. There are many ways to perform hyper-parameter searching, such as grid search, random search, and manual search [55]. Grid search is a systematic way of finding the best hyperparameter setting by training models using all possible settings automatically, which are determined by the preselected ranges of values of the hyperparameters. In this study, we incorporated grid search into our program by using the grid search procedure provided in the scikit-learn Python package [56]. We conducted grid search for all ensemble learning methods and all comparison methods included in this study.

With researchers devoting, grid search has been approved in various fields, as it has been found to improve the machine learning prediction model performance, such as the prediction of HIV/AIDS [57], text classification [58], and short-term PV power forecasting [37]. In this research, we proved that grid search can also improve the performance of ensemble learning in ubiquitination-site prediction tasks.

### Performance Metrices, 5-fold Cross-validation, and Statistical Test

We performed grid search and recorded 64 different output values for each of the models trained through an output format that we designed. Contained within the output data is information about the computer system used, computation time, and measures for model performance. For a given binary diagnostic test, a receiver operator characteristic (ROC) curve plots the true positive rate against the false positive rate for all possible cutoff values [59]. The area under curve (AUC) measures the discrimination performance of a model. We conducted 5-fold cross-validation to train and evaluate each model in a grid search. The entire dataset was split into a train-validation set, containing 80 percent of the cases, and an independent test set, containing the remaining 20 percent. We then performed a 5-fold cross validation by dividing evenly the train-validation set into 5 portions. The division was mostly done randomly except that each portion had approximately 20% of the positive cases and 20% of the negative cases to ensure that it was a representative fraction of the dataset. Training and testing were repeated five times. Each time, a unique portion was used as the validation set to test the model learned from the training set, which combined the remaining four portions. Training and testing AUCs were reported. The average training and testing AUC over all five times were also derived and reported. The best-performing set of hyperparameter values was chosen based on the highest mean test AUC. The best model would be the one refitted from the entire train-validation set using the best-performing set of hyperparameters values. We used this procedure for all methods.

We conducted statistical testing to further evaluate the prediction performance of the ensemble methods. We did both two-sided and one-sided tests. Wilcoxon rank-sum tests were performed to rank the 4 ensemble methods in terms of their prediction performance. We also compared the ensemble learning methods with traditional machine learning methods, again using both two-sided and one-sided Wilcoxon rank-sum tests. With the two-sided testing, we try to determine whether there is a significant difference between the comparison methods in terms of their prediction performance, and if there is, we would try to determine which method is better with the one-sided test.

## RESULTS

We compared the prediction performance of the best models which were selected by grid search in different aspects. The results are shown in Table 3–8 and Figure 2–5. Table 3 shows the side-by-side comparisons of the validation AUCs of the best performing models of all four ensemble methods for each of the six PCP datasets. Also included in Table 3 are the average and maximum validation AUCs for each method over all datasets and for each dataset over all methods. Based on Table 3, on average of all datasets, the ranking of the four ensemble methods is as follows: XGB (1st, AUC 0.721), ADB (2nd, 0.720), RF (3rd, 0.706), and GB (4th, 0.689). Table 4 contains the running time information and number of models trained via grid search in this study. The hyperparameters settings for the best models identified by grid searches shown in Appendix Table 1.

**Table 3.** The validation AUCs of the best-performing models recommended by grid search

|  | PCP-1 | PCP-2 | PCP-3 | PCP-4 | PCP-5 | PCP-6 | Average |
| --- | --- | --- | --- | --- | --- | --- | --- |
| RF | 0.704 | 0.665 | 0.698 | 0.719 | 0.732 | 0.718 | 0.706 |
| ADB | **0.767** | 0.671 | 0.707 | 0.708 | **0.745** | **0.723** | 0.720 |
| GB | 0.693 | 0.682 | 0.711 | 0.704 | 0.691 | 0.650 | 0.689 |
| XGB | 0.747 | **0.683** | **0.716** | **0.746** | 0.742 | 0.690 | **0.721** |
| Average | 0.728 | 0.675 | 0.708 | 0.719 | 0.728 | 0.695 | 0.709 |
| Maximum | 0.767 | 0.683 | 0.716 | 0.746 | 0.745 | 0.723 | 0.721 |

**Table 4.**
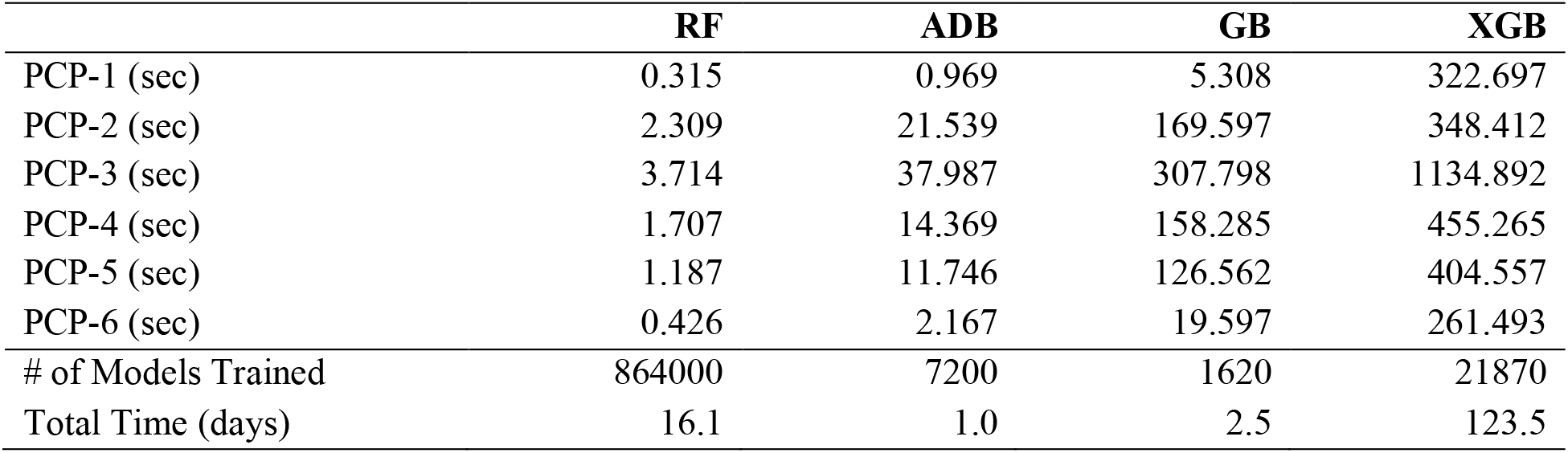
Experiment time per model, number of models trained, and total running time

**Table 5.** Two-sided Wilcoxon rank-sum test

| two-tailed | W | p-value | 95% confidence |
| --- | --- | --- | --- |
| RF vs ADB | 12 | 0.3939 | [-0.049, 0.025] |
| RF vs GB | 27.5 | 0.1488 | [-0.013, 0.048] |
| RF vs XGB | 13 | 0.4848 | [-0.048, 0.028] |
| ADB vs GB | 29 | 0.09307 | [-0.004, 0.073] |
| ADB vs XGB | 17 | 0.9372 | [-0.039, 0.040] |
| GB vs XGB | 8 | 0.132 | [-0.064, 0.008] |

**Table 6.** One-sided Wilcoxon rank-sum test

| Greater | W | p-value | 95% confidence |
| --- | --- | --- | --- |
| RF vs ADB | 12 | 0.8452 | [-0.047, inf] |
| RF vs GB | 27.5 | **0.07441** | [-0.006, inf] |
| RF vs XGB | 13 | 0.803 | [-0.043, inf] |
| ADB vs RF | 24 | 0.197 | [-0.012, inf] |
| ADB vs GB | 29 | **0.04654** | [0.003, inf] |
| ADB vs XGB | 17 | 0.5909 | [-0.038, inf] |
| GB vs RF | 8.5 | 0.9457 | [-0.039, inf] |
| GB vs ADB | 7 | 0.9675 | [-0.063, inf] |
| GB vs XGB | 8 | 0.9535 | [-0.056, inf] |
| XGB vs RF | 23 | 0.2424 | [-0.016, inf] |
| XGB vs ADB | 19 | 0.4686 | [-0.029, inf] |
| XGB vs GB | 28 | **0.06602** | [-0.001, inf] |

**Table 7.**
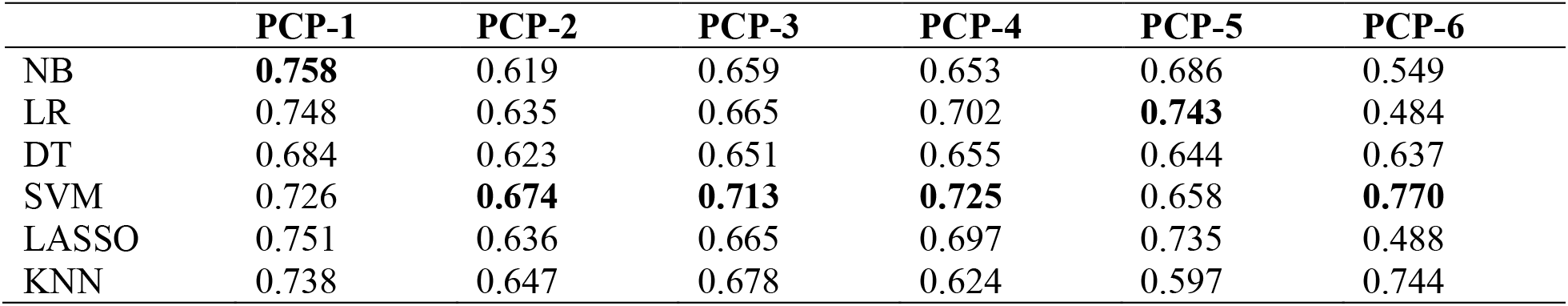
Validation score of machine learning methods

**Table 8.** One-sided Wilcoxon rank-sum test on ensemble methods vs ML methods

| **Greater** | **W** | **p-value** | **95% confidence** |
| --- | --- | --- | --- |
| RF vs NB | 29 | 0.04654 | [0.006, inf] |
| RF vs DT | 35 | 0.002165 | [0.034, inf] |
| RF vs KNN | 23 | 0.2424 | [-0.025, inf] |
| ADB vs NB | 30 | 0.03247 | [0.012, inf] |
| ADB vs DT | 35 | 0.002165 | [0.039, inf] |
| ADB vs KNN | 27 | 0.08983 | [-0.015, inf] |
| GB vs NB | 26 | 0.1201 | [-0.009, inf] |
| GB vs DT | 32 | 0.01299 | [0.013, inf] |
| GB vs KNN | 23 | 0.2424 | [-0.045, inf] |
| XGB vs NB | 29 | 0.04654 | [0.004, inf] |
| XGB vs DT | 35 | 0.002165 | [0.039, inf] |
| XGB vs KNN | 29 | 0.04643 | [0.002, inf] |

**Figure 2.**
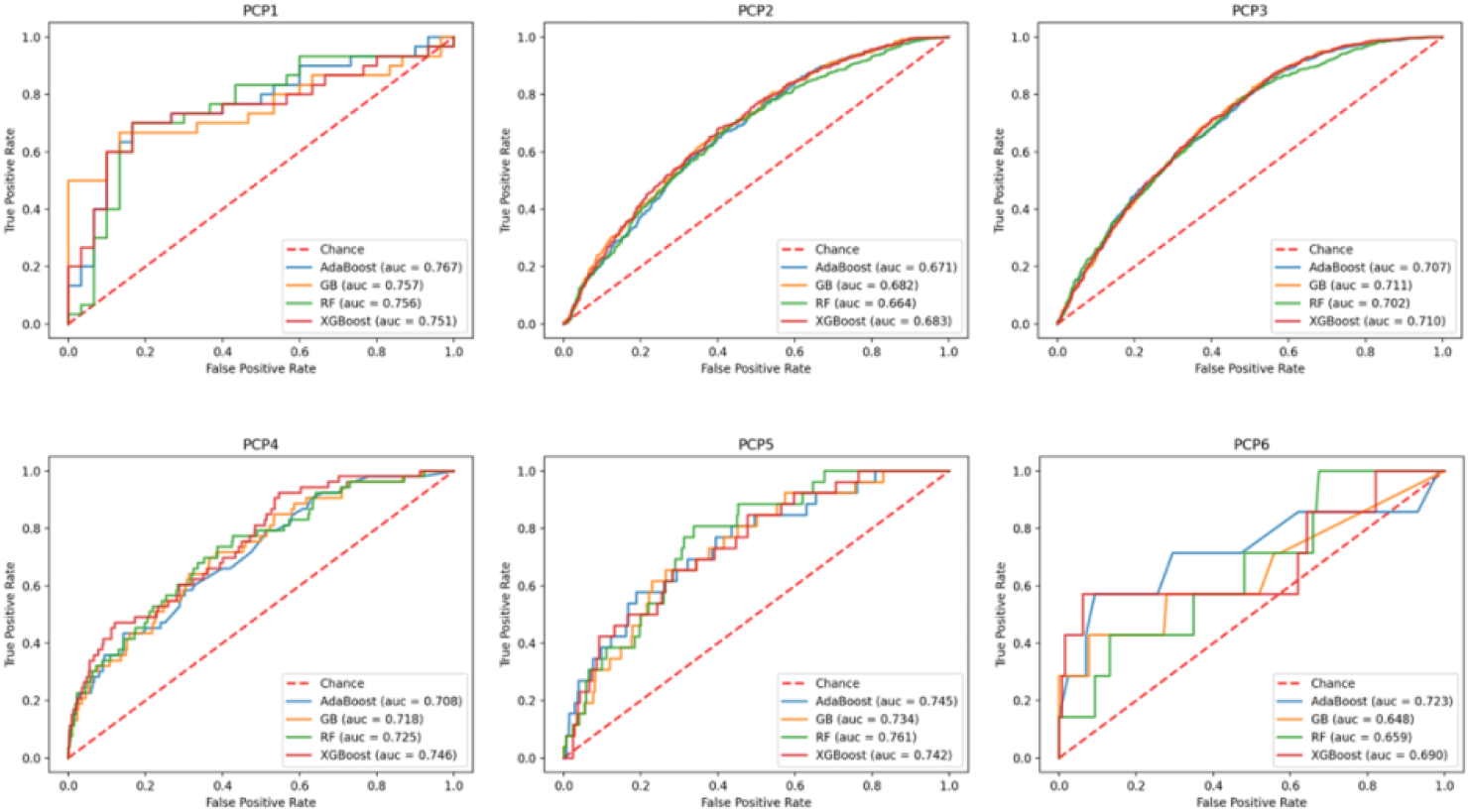
ROC curves of different best models selected by grid search

**Figure 3.**
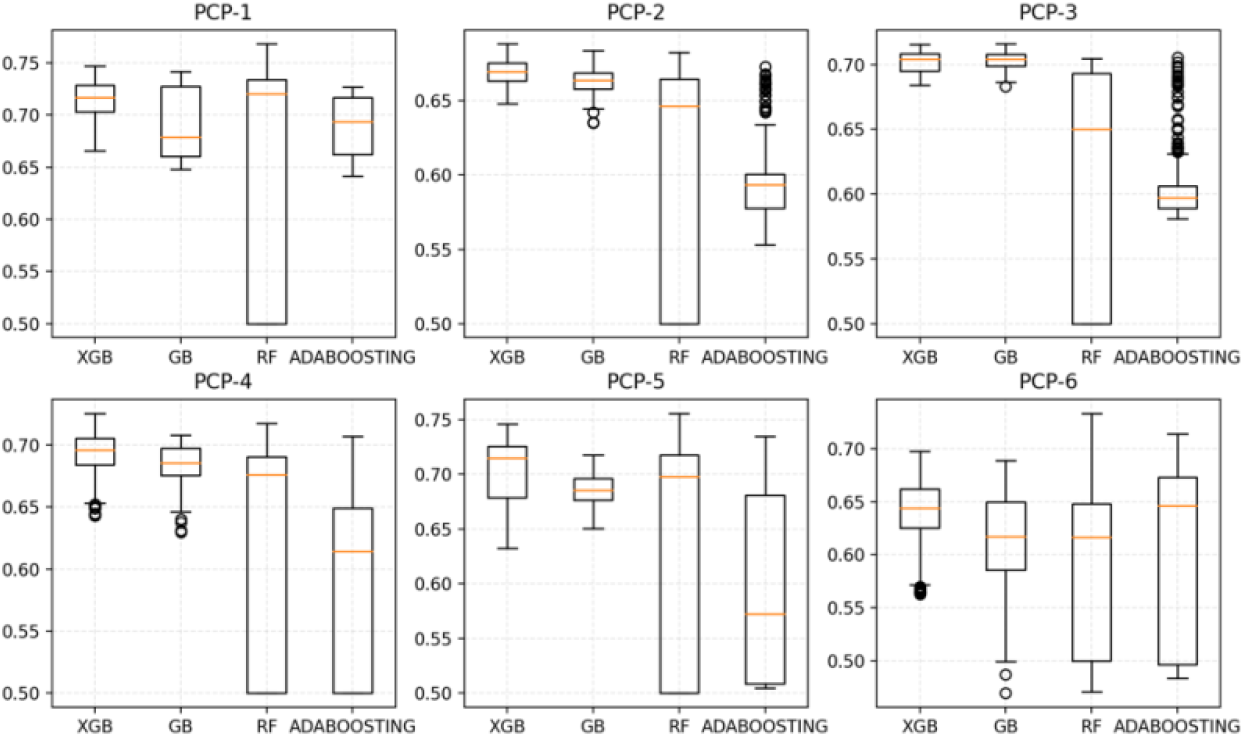
Mean test AUCs of all methods for each of the datasets

**Figure 4.**
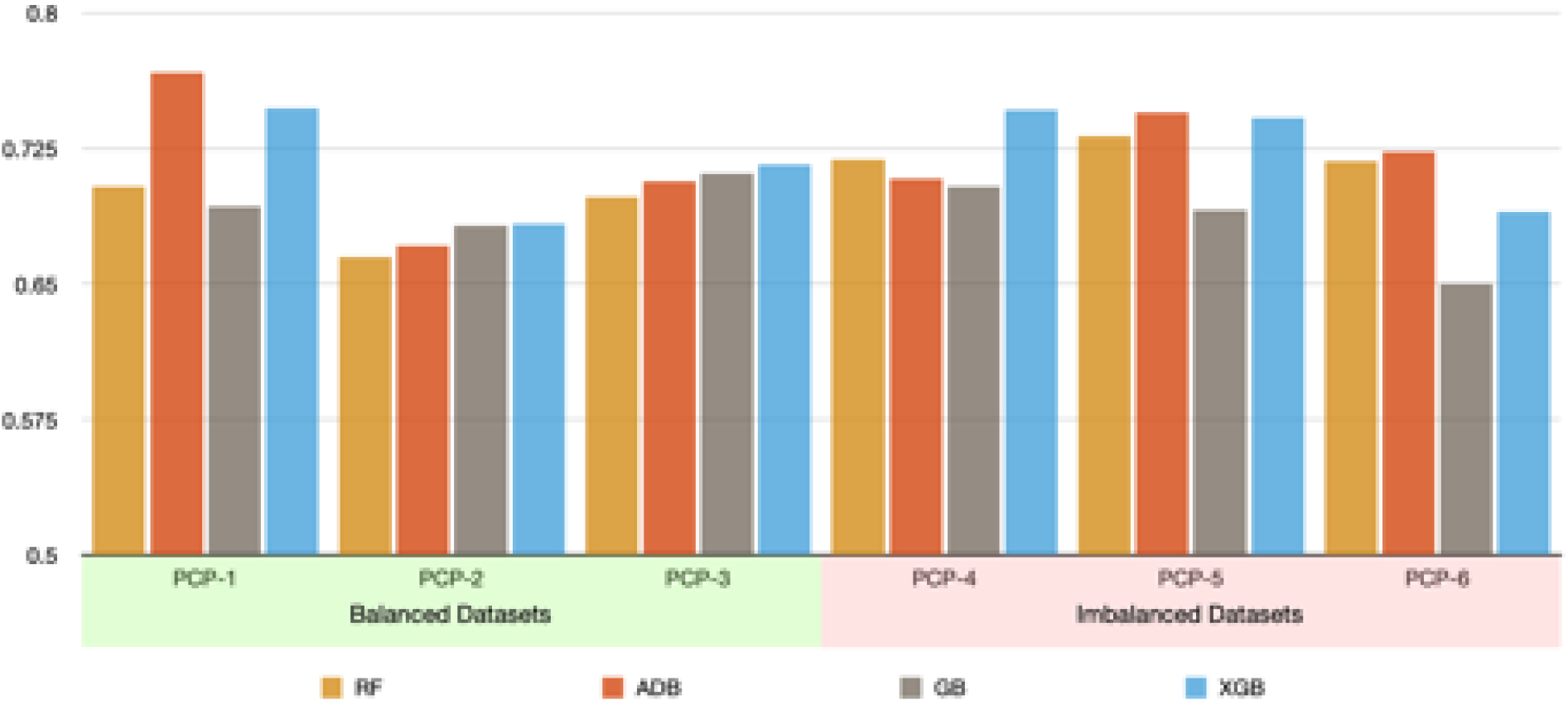
Performance comparison of balanced vs. imbalanced datasets

**Figure 5.**
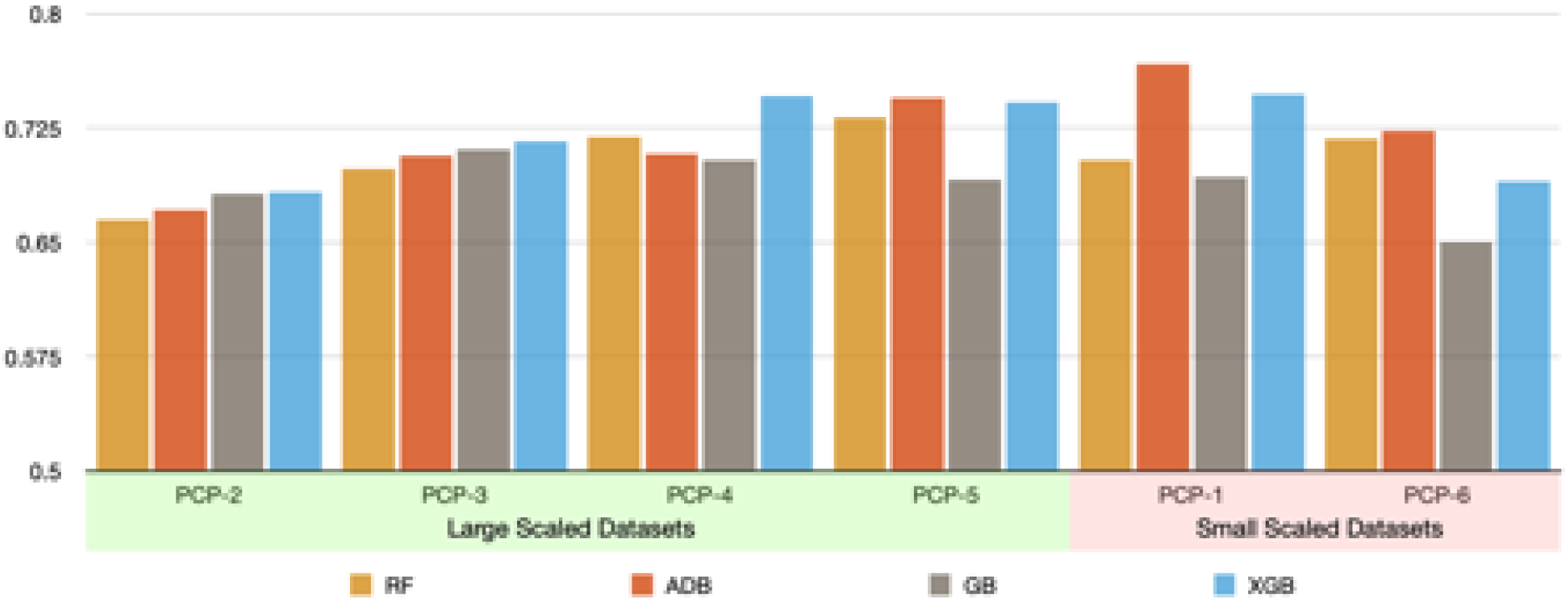
Performance comparison of large-scale and small-scale datasets

Figure 2 shows the side-by-side comparisons of the ROC curves of the four ensemble methods. It contains 6 sub-figures, one for each of the 6 PCP datasets. Based on Figure 2, all methods performed comparably for the datasets PCP-1, and PCP-2. The performance differs somewhat for datasets PCP-1, PCP-4, and PCP-5, and differs the most for dataset PCP-6. Figure 3 shows the boxplots we generated to compare side by side the prediction performance of all methods based on the mean and standard deviation of mean testing AUCs resulted from 5-fold cross-validation across all hyperparameter settings of the grid search.

As shown in Table 5, we conducted two-sided pair-wise Wilcoxson rank-sum tests among the four ensemble methods. Based on this table, at a significance level of 0.05, none of the p-values is small enough for us to reject the null hypothesis, which states that the two comparison methods perform the same. However, when we raise the significance level to 0.1, we are confident in rejecting the null hypothesis for the pair of ADB and GB. When we raise the significance level to 0.15, we are confident in rejecting the null hypothesis for the pairs of XGB and GB, and RF and GB. So based on Table 5, no evidence shows that the four methods perform differently when we allow up to 5% chance of type I error, but when we allow higher error rates, we are more confident in stating that GB performs differently from the other three methods.

Table 6 contains our one-sided pair-wise Wilcoxson rank-sum test results. Based on Table 6, at the significance level of 0.05, ADB performed better than GB. At the significance level of 0.1, we are confident that both XGB and RF perform better than GB. Both Table 5 and Table 6 show XGB, ADB, and RF do not perform differently when the type I error rate is not allowed to exceed 5%. However, based on Table 6, when we allowed the type I error rate to be as high as 25%, both XGB and ADB would perform significantly better than RF. Therefore, our statistical testing results show that XGB and ADB overall perform the same, and they both perform significantly better than RF if we allow up to 25% change for type I error. When we allow error rates up to 10%, we are confident that GB is the worst performer among all methods. Based on the p-values found in Table 5 and Table 6, we rank these 4 methods as follows: XGB≈ADB>RF>GB.

We also compared the ensemble methods with some of the non-ensemble methods including Naïve Base (NB), Logistic Regression (LR), Decision Trees (DT), Support Vector Machine (SVM), LASSO, and k-Nearest Neighbor (KNN). Table 7 shows the validation AUC results we obtained for the six non-ensemble methods using grid search. Appendix Table 2 shows all the 24 two-sided pair-wise Wilcoxon rank-sum test results between the 4 ensemble methods and the 6 non-ensemble methods. Based on this table, at the significance level of 0.1, we found the following pairs performed significantly differently (bold): RF vs NB, RF vs DT, ADB vs NB, ADB vs DT, GB v DT, XGB vs NB, XGB vs DT, and XGB vs KNN. When a two-sided pairwise test showed significant results, we further conducted a one-sided (greater than) pair-wise Wilcoxon rank-sum test for the corresponding pair. These results are contained in Table 8.

To further understand a dataset influences ensemble learning, we grouped the 6 datasets in terms of both data imbalance and the size of the data. Based on data balance, we created two groups: PCP-1, PCP-2, and PCP-3 belong to the balanced group, and PCP-4, PCP-5, and PCP-6 belong to the imbalanced group. The comparison between these two groups is shown in Figure 4. Based on the size of the data, we created large-scaled and small-scale groups. The large-scale group contains PCP-2, PCP-3, PCP-4 and PCP-5, and the small-scale group contains PCP-1 and PCP-6. The comparison between these two groups is shown in Figure 5.

## DISCUSSION

As shown in the results section, we compared the prediction performance of the four ensemble methods. GB comes with the worst validation AUC (with PCP-6) among all methods and datasets.Our statistical tests also show that GB tends to perform worst among all methods. This is perhaps because GB is one of the earlier and therefore less mature boosting methods. As shown in Figure 4 and 5, we investigated the possible effect of the dataset itself on the performance of a method. We notice that GB tends to perform comparably with other ensemble methods for large and balanced datasets such as PCP-2 and PCP-3. For small-scale and/or imbalanced datasets such as PCP-1 and PCP-6, GB tends to perform the worst. This may indicate that GB is less adaptive to data scarcity and imbalance. We notice that GB has a smaller number of hyperparameters than the later boosting methods such as ADB and XGB. It may be in the lack of hyperparameters that can be tuned to adapt to unfavorable data conditions.

Our results also show that ADB and XGB are overall very comparable in terms of their prediction performance, and each leads three of the six datasets; that is, ADB performs the best with PCP-1, PCP-5, and PCP-6; while XGB performs the best with PCP-2, PCP-3, and PCP-4. We also found that both ADB and XGB perform better than RF in most cases. As shown by Figure 3, XGB and GB can provide much stable results in all datasets regardless of what hyperparameter values are given, while RF’s interquartile ranges are the largest across all datasets, which can range from 0.5 up to 0.75. ADB has a mixed situation: it has a stabler performance for dataset PCP-1, PCP-2, and PCP-3 than it does for dataset PCP-4 through PCP-6.

We compared the ensemble methods with some of the non-ensemble methods including NB, LR, DT, SVM, LASSO, and KNN. Based on results, XGB performs significantly better than KNN; XGB, ADB, and RF perform significantly better than both NB and DT, and all ensemble methods perform better than DT. So, XGB performs better than most of the non-ensemble methods that we included in the study, ADB and RF performs better than half of the non-ensemble methods included. Even GB, the ensemble method that did the worst in this study, performed significantly better than DT. However, on the other hand, we notice that the non-ensemble methods SVM and LASSO, when using grid search, perform comparably with all the ensemble methods. The non-ensemble method KNN performs comparably with three of the four ensemble methods.

We also looked into how these ensemble methods perform differently in terms of a specific dataset by paying attention to data scarcity and imbalance issues. We notice from Table 3 that ADB, even though ranks No. 2 on average, can achieve a validation AUC as high as 0.767 (with PCP-1), which is the highest score we ever obtained using these PCP datasets. In terms of PCP-6, a small and imbalanced dataset, again ADB and XGB outperformed the other two, and ADB reached a validation AUC of 0.723, which is the highest we observed for PCP-6.

Based on Figure 4 and 5, XGB outperforms ADB for all datasets except for datasets PCP-1 and PCP-6. Note that PCP-1 and PCP-6 are the two small-scale datasets; this may indicate that ADB tends to handle small-scale datasets better than XGB. RF performs worse than XGB for all the datasets except for PCP-6, which is both a small-scale and imbalanced dataset. This may indicate that RF tends to handle this type of dataset better than XGB. Based on Figure 4, we also notice that RF tends to perform better with the imbalanced datasets than it does with the balanced datasets.

Grid search helps greatly to identify the best hyperparameter setting for each method by training a large number of models. Based on Table 5, RF is the method for which the largest number of models are trained in grid search. This is because RF has more hyperparameters than other methods. However, as shown in Table 5, it takes the least amount of time to train a model with RF among all methods. RF ends up being No. 2 in terms of the total amount of time it takes to run the grid search. ADB takes the least amount of time for grid search, because it is relative fast to train a ADB model and the total number of ADB models trained in grid search is way less than RF. XGB is the bottleneck in terms of grid search running time among all methods. This is because its unit model training times are the longest and the number of models trained in grid search ranks at No. 2 among all methods.

## CONCLUSIONS

Based on this study, ensemble learning is a promising way of improving ubiquitination-site prediction using PCP data. We find that all four ensemble methods significantly outperform one or more non-ensemble methods included in this study. XGB outperforms 3 out of the 6 non-ensemble methods that we included; ADB and RF both outperform 2 of the 6 non-ensemble methods included; GB performs better than one non-ensemble method. Comparing the four ensemble methods internally, GB performs the worst; XGB and ADB are very comparable in terms of prediction. But ADB beats XGB by far in terms of both the unit model training time and total running time (ADB’s 1 day vs XGB’s 123 days). Both XGB and ADB tend to do better than RF in terms of prediction, but RF has the shortest unit model training time out of the three. In addition, we notice that ADB tends to outperform XGB when dealing with small-scale datasets, and RF can outperform either ADB or XGB when data are less balanced. Interestingly, we find that SVM, LR, and LASSO, three of the non-ensemble methods included, perform comparably with all the ensemble methods.

## KEY POINTS

- All four ensemble methods significantly outperform one or more non-ensemble methods included in this study. Ensemble learning is a promising approach to improve ubiquitination-site prediction.
- AdaBoosting and XGBoosting tend to outperform other Ensemble methods. They are very comparable in ubiquitination-site prediction performance, but AdaBoosting needs far less time to train a model.
- AdaBoosting tends to perform better than XGBoosting when dealing with small-scale PCP datasets.
- Random Forest can outperform AdaBoosting or XGBoosting with imbalanced datasets.
- With grid search, the non-ensemble methods SVM, LR and LASSO and the ensemble methods perform comparably in ubiquitination-site prediction tasks.

## AUTHORS’ CONTRIBUTIONS

Conceptualization, XJ; methodology, XM and XJ; software, XJ; validation, XM and XJ; formal analysis, XM and XJ; investigation, XJ; resources, XJ; data curation, N/A; writing-original draft preparation, XM and XJ; writing-review and editing, XM and XJ; visualization, N/A; supervision, XJ; project administration, XJ; funding acquisition, XJ. All authors have read and agreed to the published version of the manuscript.

## DATA AVAILABILITY

The datasets used and/or analyzed during the current study are existing datasets previously published at https://bmcbioinformatics.biomedcentral.com/articles/10.1186/s12859-016-0959-z

## APPENDIX

**Appendix Table 1.**
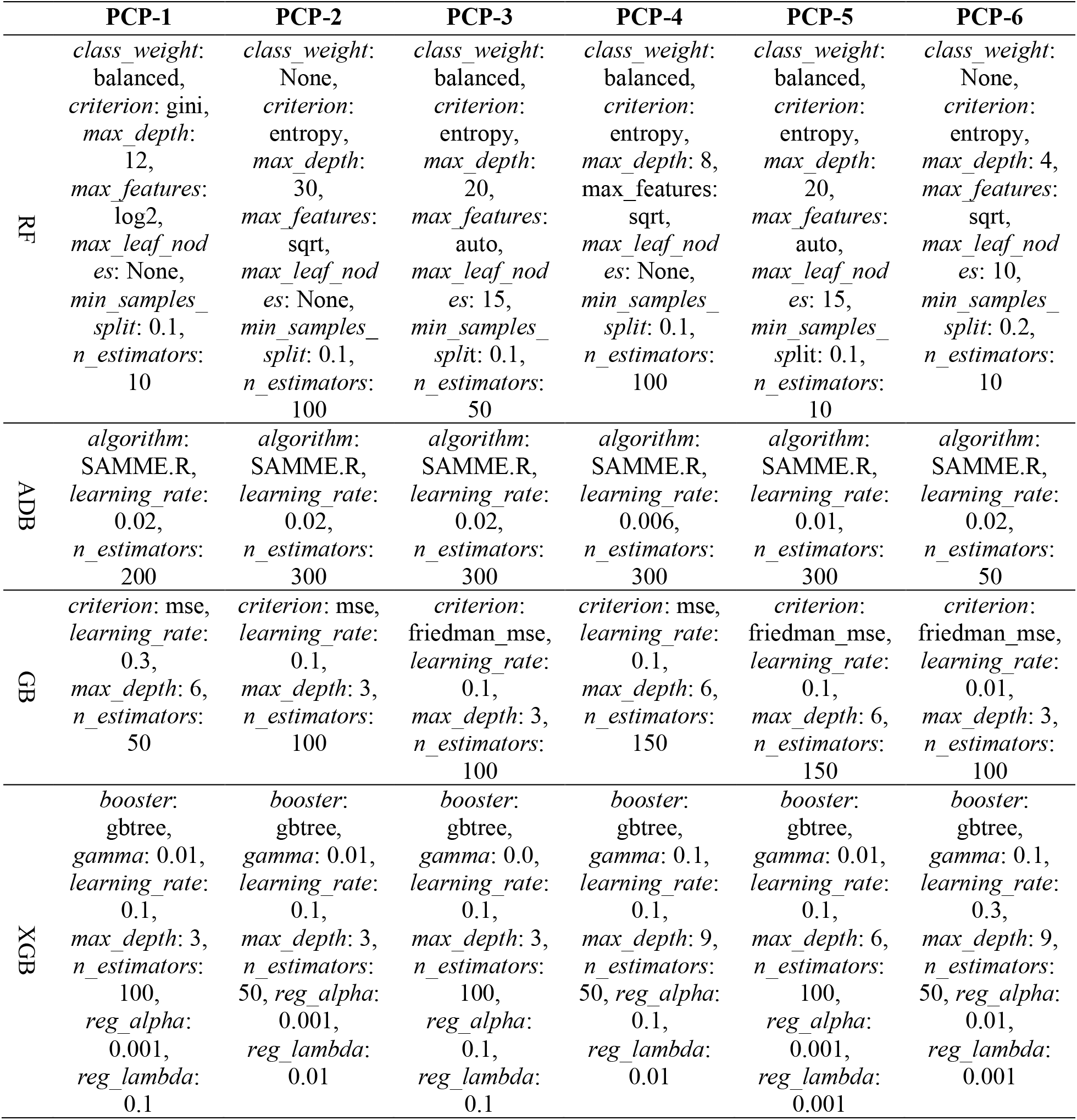
Hyperparameter values of the best-performing models learned from 6 PCP datasets.

|  | PCP-1 | PCP-2 | PCP-3 | PCP-4 | PCP-5 | PCP-6 |
| --- | --- | --- | --- | --- | --- | --- |
| RF | *class_weight*:<br>balanced,<br>*criterion*: gini,<br>*max_depth*:<br>12,<br>*max_features*:<br>log2,<br>*max_leaf_nod*<br>*es*: None,<br>*min_samples_*<br>*split*: 0.1,<br>*n_estimators*:<br>10 | *class_weight*:<br>None,<br>*criterion*:<br>entropy,<br>*max_depth*:<br>30,<br>*max_features*:<br>sqrt,<br>*max_leaf_nod*<br>*es*: None,<br>*min_samples_*<br>*split*: 0.1,<br>*n_estimators*:<br>100 | *class_weight*:<br>balanced,<br>*criterion*:<br>entropy,<br>*max_depth*:<br>20,<br>*max_features*:<br>auto,<br>*max_leaf_nod*<br>*es*: 15,<br>*min_samples_*<br>*split*: 0.1,<br>*n_estimators*:<br>50 | *class_weight*:<br>balanced,<br>*criterion*:<br>entropy,<br>*max_depth*: 8,<br>*max_features*:<br>sqrt,<br>*max_leaf_nod*<br>*es*: None,<br>*min_samples_*<br>*split*: 0.1,<br>*n_estimators*:<br>100 | *class_weight*:<br>balanced,<br>*criterion*:<br>entropy,<br>*max_depth*:<br>20,<br>*max_features*:<br>auto,<br>*max_leaf_nod*<br>*es*: 15,<br>*min_samples_*<br>*split*: 0.1,<br>*n_estimators*:<br>10 | *class_weight*:<br>None,<br>*criterion*:<br>entropy,<br>*max_depth*: 4,<br>*max_features*:<br>sqrt,<br>*max_leaf_nod*<br>*es*: 10,<br>*min_samples_*<br>*split*: 0.2,<br>*n_estimators*:<br>10 |
| ADB | *algorithm*:<br>SAMME.R,<br>*learning_rate*:<br>0.02,<br>*n_estimators*:<br>200 | *algorithm*:<br>SAMME.R,<br>*learning_rate*:<br>0.02,<br>*n_estimators*:<br>300 | *algorithm*:<br>SAMME.R,<br>*learning_rate*:<br>0.02,<br>*n_estimators*:<br>300 | *algorithm*:<br>SAMME.R,<br>*learning_rate*:<br>0.006,<br>*n_estimators*:<br>300 | *algorithm*:<br>SAMME.R,<br>*learning_rate*:<br>0.01,<br>*n_estimators*:<br>300 | *algorithm*:<br>SAMME.R,<br>*learning_rate*:<br>0.02,<br>*n_estimators*:<br>50 |
| GB | *criterion*: mse,<br>*learning_rate*:<br>0.3,<br>*max_depth*: 6,<br>*n_estimators*:<br>50 | *criterion*: mse,<br>*learning_rate*:<br>0.1,<br>*max_depth*: 3,<br>*n_estimators*:<br>100 | *criterion*:<br>friedman_mse,<br>*learning_rate*:<br>0.1,<br>*max_depth*: 3,<br>*n_estimators*:<br>100 | *criterion*: mse,<br>*learning_rate*:<br>0.1,<br>*max_depth*: 6,<br>*n_estimators*:<br>150 | *criterion*:<br>friedman_mse,<br>*learning_rate*:<br>0.1,<br>*max_depth*: 6,<br>*n_estimators*:<br>150 | *criterion*:<br>friedman_mse,<br>*learning_rate*:<br>0.01,<br>*max_depth*: 3,<br>*n_estimators*:<br>100 |
| XGB | *booster*:<br>gbtree,<br>*gamma*: 0.01,<br>*learning_rate*:<br>0.1,<br>*max_depth*: 3,<br>*n_estimators*:<br>100,<br>*reg_alpha*:<br>0.001,<br>*reg_lambda*:<br>0.1 | *booster*:<br>gbtree,<br>*gamma*: 0.01,<br>*learning_rate*:<br>0.1,<br>*max_depth*: 3,<br>*n_estimators*:<br>50, *reg_alpha*:<br>0.001,<br>*reg_lambda*:<br>0.01 | *booster*:<br>gbtree,<br>*gamma*: 0.0,<br>*learning_rate*:<br>0.1,<br>*max_depth*: 3,<br>*n_estimators*:<br>100,<br>*reg_alpha*:<br>0.1,<br>*reg_lambda*:<br>0.1 | *booster*:<br>gbtree,<br>*gamma*: 0.1,<br>*learning_rate*:<br>0.1,<br>*max_depth*: 9,<br>*n_estimators*:<br>50, *reg_alpha*:<br>0.1,<br>*reg_lambda*:<br>0.01 | *booster*:<br>gbtree,<br>*gamma*: 0.01,<br>*learning_rate*:<br>0.1,<br>*max_depth*: 6,<br>*n_estimators*:<br>100,<br>*reg_alpha*:<br>0.001,<br>*reg_lambda*:<br>0.001 | *booster*:<br>gbtree,<br>*gamma*: 0.1,<br>*learning_rate*:<br>0.3,<br>*max_depth*: 9,<br>*n_estimators*:<br>50, *reg_alpha*:<br>0.01,<br>*reg_lambda*:<br>0.001 |

**Appendix Table 2.**
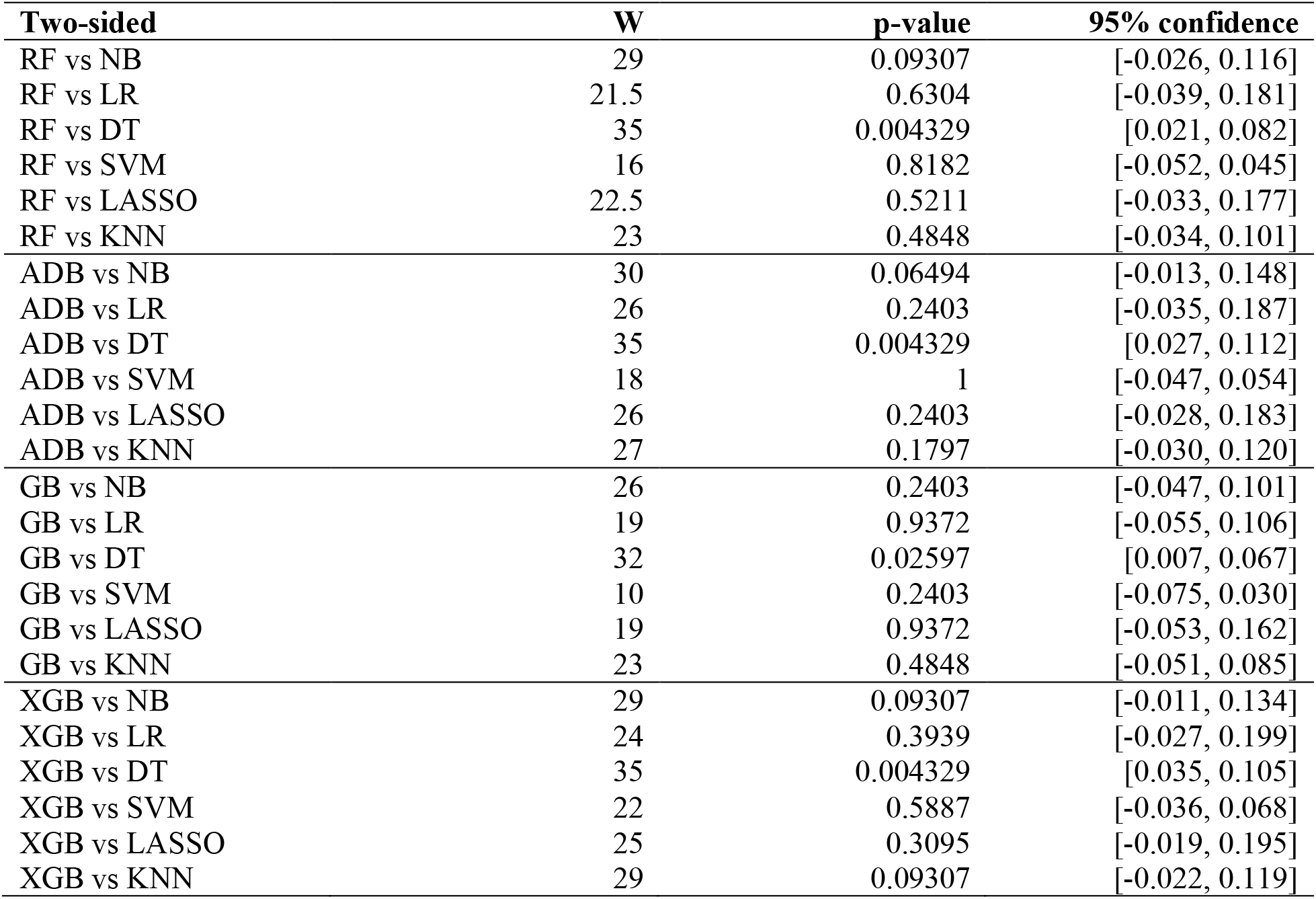
Two-sided Wilcoxon rank-Sum test on ensemble methods vs ML methods.

